# Understanding Strategic Motor Learning as a Process of Hypothesis Testing

**DOI:** 10.64898/2026.09.05.749306

**Authors:** Anjuli Niyogi, Elizabeth Cisneros, Wei Ding, Kelsey Allen, Saurabh Vyas, Richard Ivry, Jonathan Tsay

**Affiliations:** Department of Psychology, Carnegie Mellon University; Department of Psychology, University of California, Berkeley; Helen Wills Neuroscience Institute, University of California, Berkeley; Department of Psychological and Cognitive Sciences, Tsinghua University; Department of Computer Science at the University of British Columbia; Department of Psychology at the University of British Columbia; Neuroscience Institute, Carnegie Mellon University; Biomedical Engineering, Carnegie Mellon University

## Abstract

Multiple learning processes contribute to successful goal-directed actions under changing physiological states, biomechanical constraints, and environmental contexts. Among these, explicit strategies enable us to discover new movement patterns when existing ones no longer achieve the desired outcome. Yet, how strategies are discovered during motor learning remains unknown. To address this, we developed a novel behavioral paradigm that isolates strategy discovery in response to a range of visuomotor perturbations. This approach revealed that strategy discovery unfolds through an initial period of systematic exploration across multiple candidate strategies, followed by an “aha” moment in which behavior converges on a stable solution. To account for these dynamics, we developed a computational model based on hypothesis testing in which learners generate and evaluate visuomotor rules to counteract the perturbation. This hypothesis testing model outperformed a range of alternative accounts, including gradual error reduction, win-stay lose-shift learning, and sudden one-shot insight. Together, these findings identify hypothesis testing as a computational mechanism by which humans discover new strategies during motor learning.

## Introduction

Picture an amateur golfer who has hit an errant drive and now finds their ball resting just in front of a large tree. The situation is unlike any they have faced before: 200 yards to the green but not enough space to take a full swing. Success here does not come from *refining* a familiar technique, but from *discovering* a new one altogether. This ability to generate effective actions in novel situations is a defining feature of human motor learning, arising from the interplay of multiple learning processes – some implicit and automatic, others explicit and strategic (Krakauer et al., 2019; Redding & Wallace, 1996; Tsay et al., 2022, 2024).

Our understanding of motor learning has been shaped almost entirely by models of implicit motor learning. In these models, the motor system gradually minimizes sensory prediction errors—the discrepancy between intended and observed sensory outcomes—to iteratively optimize movement toward a single correct solution (Kim et al., 2021; Krakauer et al., 2019; Shadmehr et al., 2010; Tsay et al., 2022; Tseng et al., 2007). As a result, motor learning has been viewed as the gradual refinement of an existing movement.

Yet many motor behaviors require more than refining an existing movement—they require discovering an entirely new strategy (e.g., inventing the Fosbury flop). This discovery problem has long been recognized, from the ‘cognitive’ stage of skill acquisition proposed by Fitts and Posner to the recent surge of interest in de novo motor learning (Fitts & Posner, 1967; Sternad, 2018; Yang et al., 2021). Surprisingly, however, the computational mechanisms that enable motor strategy discovery remain unknown.

Emerging evidence suggests that motor strategy discovery operates through fundamentally different principles than implicit motor refinement. First, rather than variability being attributed solely to sensorimotor noise, strategy use can lead to variability that clusters into multiple discrete modes of movement, consistent with learners generating and evaluating competing candidate solutions (Ding et al., 2026). Second, rather than progressively adjusting toward a single solution, strategic behavior often culminates in an “aha” moment in which performance rapidly stabilizes on one of several possible solutions (Eliopulos et al., 2026; Townsend et al., 2026). Collectively, these findings suggest that strategic motor learning emerges not through gradual error reduction, but through systematic search over possible solutions.

Here, we provide a new computational account of how people discover motor strategies during sensorimotor learning. Specifically, learners may discover movement strategies by generating, evaluating, and eliminating competing visuomotor rules linking actions to their sensory outcomes. This idea parallels theories of human problem solving, in which hypothesis testing serves as a core mechanism for uncovering hidden structure in uncertain environments (AbdelRahman et al., 2026; Bower & Trabasso, 1963; Lee et al., 2016; Piantadosi et al., 2016; Pitt et al., 2026; Restle, 1962; Simon, 1975). Consider the sequence 2, 4, 8. One might hypothesize that the pattern reflects repeated doubling and therefore predict the next value to be 16. If the next value is revealed to be 14 instead, this hypothesis is falsified, prompting the learner to evaluate alternative hypotheses. Through successive cycles of prediction and evaluation, successful learners eventually infer the underlying rule that each term increases by a progressively larger increment (+2, +4, +6, …).

Motor strategy discovery may be governed by similar principles of hypothesis testing. When confronted with an unfamiliar visuomotor perturbation, learners entertain competing visuomotor rules linking their actions to their sensory outcomes (e.g., when using an unfamiliar laptop for a podium presentation, where the mapping between hand movement and cursor motion is unexpectedly rotated, reflected, or scaled on the presentation screen compared to one’s own computer setup). These candidate rules are then translated into compensatory strategies whose predictions are evaluated against sensory feedback. By this view, discrete modes of movement reflect the evaluation of competing visuomotor rules, with the accumulation of evidence eventually giving rise to an “aha” moment in which behavior converges on an effective strategy. To test this account, we introduce the Motor Inference and Discovery (MIND) task, a novel behavioral paradigm designed to isolate strategic motor learning from other forms of motor learning (Fig. 1). Specifically, participants were instructed to counteract a hidden visuomotor perturbation (rotation or reflection) by positioning a computer cursor within a two-dimensional workspace and confirming their chosen action with a mouse click. Critically, participants could deliberate for as long as they needed before committing to an action, as the cursor remained visible until they received feedback. Unlike conventional motor learning paradigms, this design minimized the influence of sensorimotor noise and implicit motor refinement, biasing behavior toward strategy discovery (Wong et al., 2019). We developed a computational model of hypothesis testing and, in three experiments, compared its performance against alternative computational accounts.

**Figure 1.**
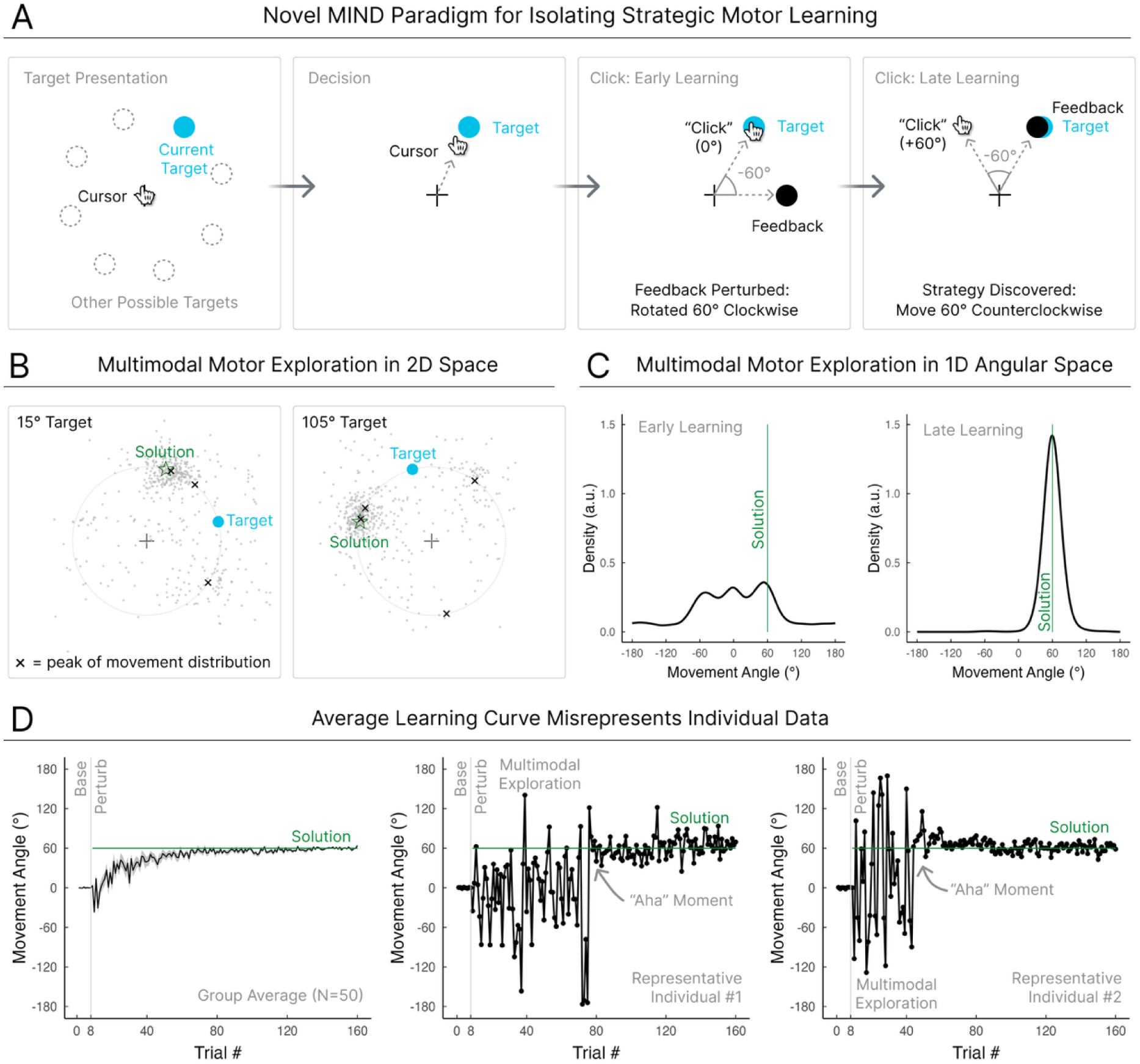
Strategic discovery is characterized by multimodal exploratory behavior that is punctuated by an “aha” moment. **(A)** Motor Inference and Discovery (MIND) task procedure. On each trial, a blue target was presented, with its position randomly selected from one of eight locations. Participants moved the cursor (hand icon) to their selected location and confirmed their choice with a mouse click. During early learning, participants explored multiple candidate solutions (including movements directly to the target location). By late learning, responses converged on the location 60° from the target, opposite the direction of the perturbation. Rotation direction was counterbalanced across participants (Experiment 1: N = 50). **(B)** Mouse click locations (gray dots) from all participants for the 15° (left) and 105° (right) targets. The blue circle denotes the target location, and the green star denotes the correct compensatory solution. Gaussian mixture model means are denoted by “x” symbols. **(C)** Distribution of movement angles relative to the target during early (left) and late learning (right), pooled across targets. **(D)** Group and individual learning trajectories. Group mean movement angle (solid line) and ± 1 SEM (shaded) are shown on the left. Representative learning trajectories from two participants are shown on the right.

## Results

### Experiment 1: Hypothesis Testing Governs Strategic Motor Learning in Response to a Visuomotor Rotation

We tested participants (N = 50) on the Motor Inference and Discovery (MIND) task, a novel behavioral paradigm designed to isolate strategic motor learning from other forms of motor learning (Fig. 1). The participants’ goal was to move to a location in the two-dimensional workspace such that a white feedback circle would be aligned with a target, a blue circle. On each trial, the participants used their computer mouse to position the cursor, displayed as a hand icon, and confirmed their choice with a mouse click. As soon as the click was recorded, the feedback circle appeared. The position of this endpoint feedback was either aligned with the cursor position (veridical feedback) or perturbed (Fig. 1). The target, computer cursor, and endpoint feedback remained visible for 2 seconds. This method, in which the computer cursor is always visible, has been shown to minimize implicit motor learning (Wong et al., 2019).

During baseline, the visual feedback was veridical, allowing participants to achieve the goal by simply moving to the target location. However, during the perturbation phase, the feedback cursor was rotated by 60° relative to the selected location (clockwise or counterclockwise, counterbalanced across participants). Consequently, successful performance required participants to infer the hidden visuomotor rule (i.e., a 60° rotation) and discover a strategy to counteract it (i.e., selecting a location 60° in the opposite direction), thereby realigning the perturbed feedback with the target (Fig. 1A). Importantly, participants were given no information about the identity of the perturbation.

At the group-level, learning appeared incremental (Fig. 1D), consistent with the view that strategy discovery is driven by gradual error reduction. During the baseline phase, participants moved directly to the target (mean movement angle ± SEM: Baseline = 0.01° ± 0.003). During the perturbation phase, the movement angle gradually shifted toward the 60° solution required to counteract the perturbation (late learning = 60.0° ± 0.01°, p < .001). This improvement was accompanied by a substantial reduction in motor variability, from broad exploration early in learning (first 16 perturbed trials; SD = 66.9° ± 3.6°) to highly consistent responses as participants converged on the correct solution late in learning (last 16 perturbed trials; SD = 10.3° ± 1.1°; Early vs Late SD: t(49) = 5.3, p < .001).

However, this group-level learning function obscured a strikingly different pattern at the level of individual learners (Fig. 1D). Rather than incrementally refining a single movement strategy toward the 60° solution, participants distributed their responses across multiple candidate strategies early in learning (Fig. 1B & Fig. S1; mean number of response clusters per target: 3.5 clusters ± 0.3; ΔBIC = 698.9). Modes emerge not only at the baseline solution (0°) and the correct solution (60°), but also at the reversal solution (180° opposite the target) and sign-flip solution (60° in the wrong direction) (Fig. 1B). The presence of these discrete, recurring movement modes is difficult to reconcile with gradual error reduction. Instead, they suggest that learners systematically explore competing visuomotor hypotheses about the hidden perturbation.

As learning progressed, individuals’ multiple response modes rapidly converged onto a single peak centered near the correct 60° solution, consistent with an “aha” moment (Fig. 1D). To quantify the timing of this transition, we applied a change-point analysis (Townsend et al., 2026) to identify when the mean and variance of each participant’s learning trajectory changed most abruptly. There was substantial variability across participants in this timepoint—the number of trials required to reach their final strategy— (range: 8–120 post-perturbation trials, median: 32). Again, this abrupt transition is difficult to reconcile with gradual error reduction and instead suggests that learners undergo discrete shifts in their beliefs about the hidden visuomotor perturbation, and the motor strategy required to counteract it.

What computational principles underlie strategy discovery during motor learning? To answer this question, we formalized and compared four competing accounts of strategy discovery, asking whether they could explain the two defining signatures observed in our data: multimodal exploration and the ‘aha’ moment. For all models, we verified that we could recover the underlying model parameters and reliably distinguish between competing models (Kovács & Lengyel, 2026; Wilson & Collins, 2019) (Fig. S2).

The first account, *gradual error reduction*, posits that learners continuously refine a single movement strategy in response to motor error—the canonical account of implicit motor learning. By this view, behavior evolves through a sequence of small, incremental corrections that gradually transform an initial baseline solution (i.e., reaching directly to the target) into the correct compensatory solution (e.g., shifting the aiming direction to 60° counterclockwise to the target to compensate for the 60° clockwise visuomotor rotation) (Donchin et al., 2003; Smith et al., 2006; Thoroughman & Shadmehr, 2000).

The second account, *moment of insight*, posits that learners discover successful strategies abruptly rather than gradually. By this view, behavior exhibits minimal exploration early in learning (variability attributed to noise in the sensorimotor system), with movements directed toward the target. Simultaneously, the participant analyzes feedback until the correct solution is identified, which triggers an abrupt transition in performance (Townsend et al., 2026).

The third account, *win-stay, lose-shift*, posits that learners discover successful strategies through stochastic exploration. Actions that produce favorable outcomes are likely to be repeated, with variability arising from sensorimotor noise, whereas unsuccessful outcomes trigger random exploration around previously rewarded actions, gradually biasing behavior toward more effective solutions (Cashaback et al., 2019; van Mastrigt et al., 2021).

For the fourth account, we developed a *hypothesis testing* model, inspired by theories of human problem solving (Piantadosi et al., 2016). Here, strategy discovery emerges through the generation and evaluation of competing visuomotor rules that link actions to their sensory outcomes. Specifically, we propose that participants search over a rule space comprising plausible perturbations—including rotations, reflections, and translations—that could account for the observed perturbation (Table S1). On each trial, an action is selected based on the participant’s current beliefs about these candidate rules. The observed sensory outcome is compared with the sensory predictions from competing rules, and the resulting prediction errors are used to update the probability assigned to each rule in parallel via Bayesian inference. As evidence accumulates, behavior transitions from systematically evaluating competing rules to exploiting the rule with the strongest evidential support.

The hypothesis testing model provided the most parsimonious account of the data (Fig. 2B). Based on the Bayesian Information Criterion (BIC), it was the best-fitting model for 90% of participants (45 of 50). Importantly, the hypothesis testing model also provided an excellent absolute account of the observed behavior at the individual level. Posterior predictive simulations faithfully reproduced the key behavioral signatures of strategic motor learning. At the individual level, the model captured both the multimodal exploration observed early in learning and the abrupt “aha” transition to the correct solution (Fig. 2A). At the group-level, the model accurately reproduced the observed distributions of movement angles early in learning, including the sign-flip solution (−60°) and the reversal solution (180°). Consistent with this qualitative agreement, the hypothesis testing model yielded the lowest Jensen-Shannon divergence from the empirical movement angle distributions (hypothesis testing: 0.26 bits; all alternative models > 1.22 bits), indicating a remarkably close match to the observed behavioral distributions. The hypothesis testing model also provided the best fit to the group-averaged learning trajectories (Fig. S3).

**Figure 2.**
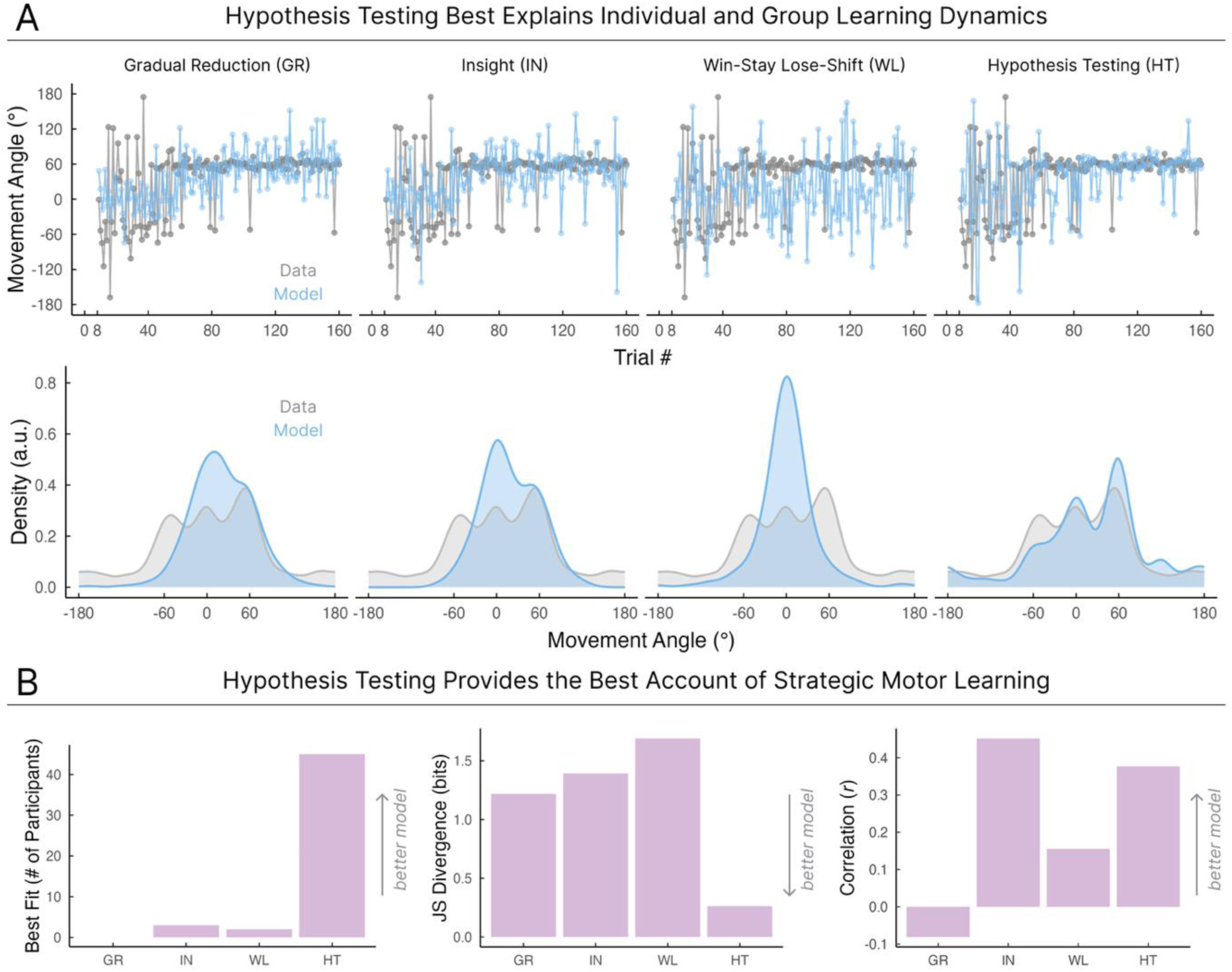
Hypothesis Testing Governs Strategic Motor Learning in Response to a Visuomotor Rotation. **(A)** Top row shows trial-by-trial movement angle data for a representative participant (grey). Each column corresponds to one of the four models, with the simulated movement angle results shown in blue. Simulations were generated using the best-fitting parameters for each model fit to each participant. Bottom row shows the distribution of movement angles during early learning. **(B)** Model comparison. Number of participants best fit by each model according to BIC (left). Jensen–Shannon distance between group-level empirical and simulated early movement angle distributions (middle; lower values indicate better agreement). Pearson’s correlation between individuals’ empirical and simulated changepoint (right; higher values indicate better agreement).

The hypothesis testing model also captured substantial between-participant variability in learning behavior. It exhibited the strongest correspondence between observed and simulated exploratory variability (standard deviation of movement angles) across individuals (*r* = .80, p < .001, 95% CI [.7, .9]; all alternative models < 0.41). Additionally, the model accurately identified when individual participants experienced their “aha” moment, exhibiting a high correlation between the observed and simulated change points (*r* = .38, p = .015, 95% CI [.08, .6]). Its performance closely matched that of the moment of insight model, which was designed specifically to capture abrupt transitions in learning (*r* = .45, p = .003) and exceeded that of the other two candidate models (win-stay lose-shift: *r* = .16, p = .33; gradual error reduction: *r* = -.08, p = .61).

In summary, these findings demonstrate that hypothesis testing provides a superior account of both aggregate behavioral signatures and individual learning dynamics, offering a unified computational account of strategic motor learning across levels of analysis.

### Experiment 2: Hypothesis Testing Governs Strategic Motor Learning in Response to a Visuomotor Reflection

Real-world motor learning requires mastering a diverse range of visuomotor transformations. For example, compensating for a constant gust of wind while golfing may require learning a visuomotor translation, operating an unfamiliar computer cursor may require adapting to a visuomotor gain, and brushing one’s teeth in a mirror may require learning a visuomotor reflection. Consequently, if hypothesis testing serves as a general computational principle of motor learning, it should account for strategy discovery across different visuomotor transformations.

To test this, Experiment 2 used the same MIND task and applied a diagonal mirror reflection as the hidden perturbation, such that visual feedback was reflected across an invisible 45° diagonal axis (N = 44). Whereas a single global transformation governs a visuomotor rotation, a mirror reflection requires target-specific compensatory movements. For some targets, successful performance requires only a modest deviation from the target, whereas for others, it requires aiming in the opposite direction in the workspace (Heirani Moghaddam et al., 2026; Telgen et al., 2014; Yang et al., 2021). Consequently, compensating for a diagonal mirror reflection stress tests the hypothesis testing account under a qualitatively different visuomotor transformation.

Despite substantial differences in rule structure, strategy discovery under mirror reflection exhibited the same signatures observed under visuomotor rotation. For direct comparison with the rotation condition, we focused our primary analyses on the 15° and 195° target locations, where the required compensatory solution was identical (60°), although all target locations showed the same pattern (Fig. S1). At the group level, participants gradually reduced their errors and converged on the solution (mean movement angle ± SEM: baseline = −0.1° ± 0.001; late learning = 49.9° ± 0.04; Early vs Late: t(43) = −7.56, p < .001), giving the appearance of gradual error reduction (Fig. 3B).

**Figure 3.**
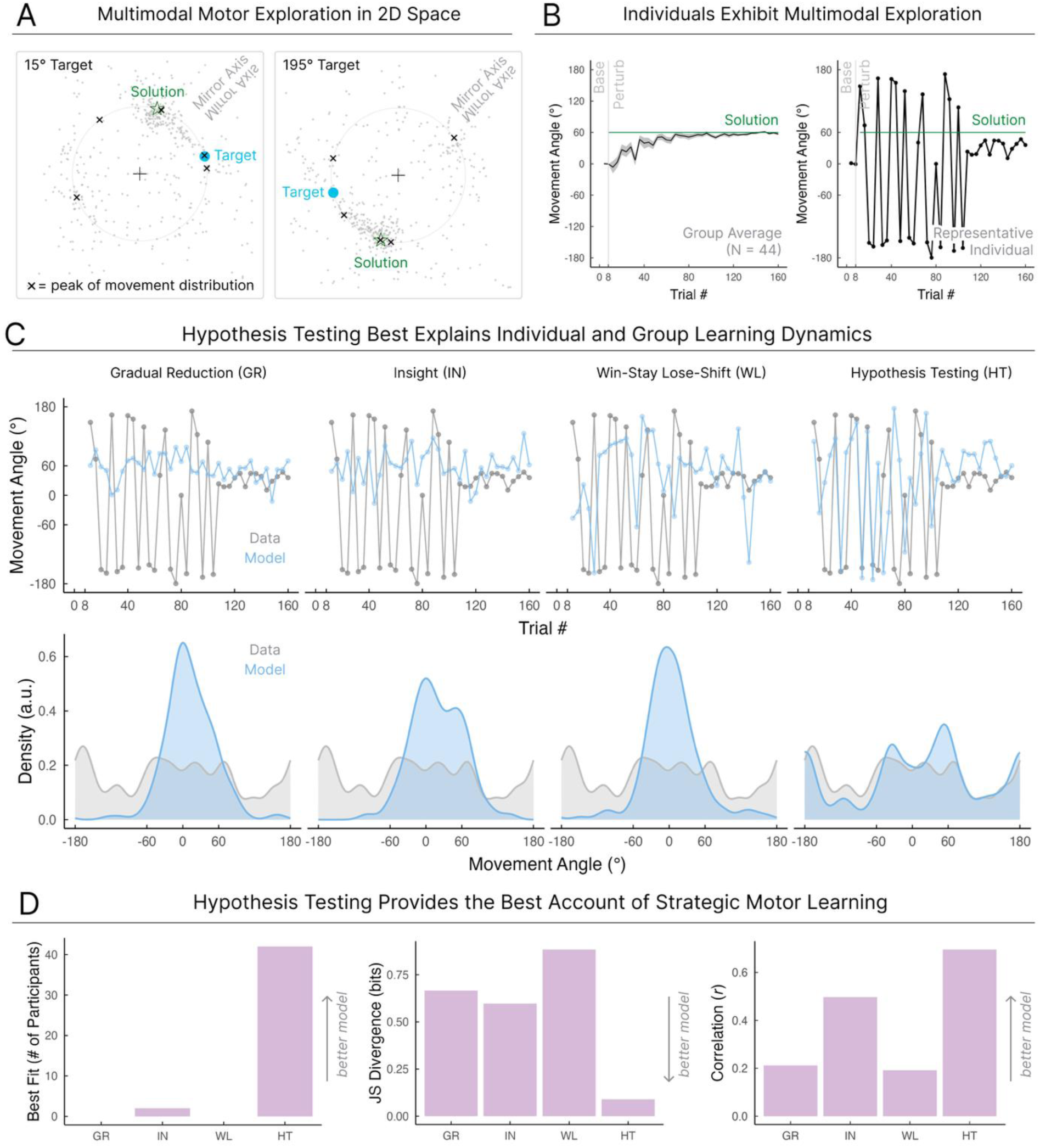
Hypothesis Testing Governs Strategic Motor Learning in Response to a Visuomotor Reflection. **(A)** Mouse click locations from all participants across the workspace for the 15° (left) and 195° (right) targets (Experiment 2: N = 44). The blue circle denotes the target location, the green star denotes the correct compensatory solution, and Gaussian mixture model angular means are denoted by “x” symbols. **(B)** Group (left) and individual (right) learning trajectories across trials. Data are shown for the 15° and 195° target locations, for which the correct compensatory solution is a 60° rotation. **(C)** Empirical behavior is shown in gray, with corresponding simulations of individual learning curves and movement-angle distributions shown in blue for each model. Simulations were generated using the best-fitting parameters for each model fit to each individual participant. Columns represent competing models. **(D)** Model comparison. Number of participants best fit by each model according to BIC (left). Jensen– Shannon divergence between group-level, target-specific empirical and simulated movement-angle distributions (middle; lower values indicate better agreement). Pearson’s correlation between individuals’ empirical and simulated changepoints (right; higher values indicate better agreement).

However, the group-level average obscured the patterns observed at the individual level (Fig. 3B): Early learning was highly multimodal, with responses clustering around five discrete candidate solutions (Fig. 3A; mean number of clusters per target: 5.6 ± 0.5; ΔBIC = 459.8). Prominent response modes across target locations corresponded to the reversal solution (180° from the target), a horizontal-axis reflection (−30°), and aiming at the target (0°). As learning progressed, these competing movement modes gave way to the correct solution, producing the characteristic “aha” moment. There was substantial variability across participants in the number of trials required to reach their final strategy (range: 10–136 post-perturbation trials, median: 51).

The hypothesis testing model again provided the most parsimonious account of the results (Fig. 3D), achieving the lowest BIC for 95% of participants (42/44). It also produced the most accurate posterior predictions (Fig. 3C), reproducing the multimodal exploratory behavior observed early in learning (lowest JS divergence: 0.09 bits; alternative models > 0.60 bits) and showed the strongest agreement between predicted and empirical movement variability (*r* = .54, p < .001, 95% CI [.4, .7]; alternative models < 0.15). The hypothesis testing model captured individual change points with accuracy superior to that of the alternative models (*r* = .70, p < .001, 95% CI [.5, .8]; alternative models < 0.5). Together, these findings suggest that strategic motor learning across diverse perturbation structures may be governed by a common computational principle: hypothesis testing.

### Experiment 3: Hypothesis Testing Explains Why Different Learners Discover Different Solutions

Consider the sequence 2, 4, 8: one person may infer that the next values are 16, 32, 64 by applying a ×2 rule, whereas another may infer 14, 22, 32 by applying a +2-addend increment rule (+6, +8, +10). Because the observed sequence is consistent with both hypotheses, the available evidence does not uniquely determine the underlying rule. We propose that this same principle extends to motor learning: when sensory evidence is ambiguous, *different* learners may converge on *different* motor strategies. This prediction arises naturally from the hypothesis testing model because learners evaluate *multiple* candidate solutions. In contrast, the alternative models considered here are typically formulated to learn a single solution and therefore, in their canonical form, do not predict which of multiple solutions different learners will discover (Piantadosi et al., 2016). Accounting for such solution diversity within these conventional models would require additional assumptions.

We tested the prediction that ambiguous sensory evidence can lead *different* learners to discover *different* motor strategies by reanalyzing a previously collected dataset from our laboratory (Ding et al., 2026). Participants (N = 54) learned a 60° clockwise visuomotor rotation with delayed endpoint feedback—a manipulation similar to the veridical feedback of the mouse-click location used in Exps. 1–2—known to minimize implicit motor learning and thereby bias behavior toward strategy use (Fig. 4A) (Brudner et al., 2016). Training was restricted to just two targets (the 80° and 100° targets). Critically, within this narrow region of the workspace, the available sensory evidence (the resulting feedback position) is consistent with at least two visuomotor rules: a global 60° clockwise rotation rule and a shift “rightward” rule (Fig. 4B). Although both visuomotor rules yield successful compensation at the trained targets, the hypothesis testing account predicts that participants may arrive at different explanations for the perturbation.

**Figure 4.**
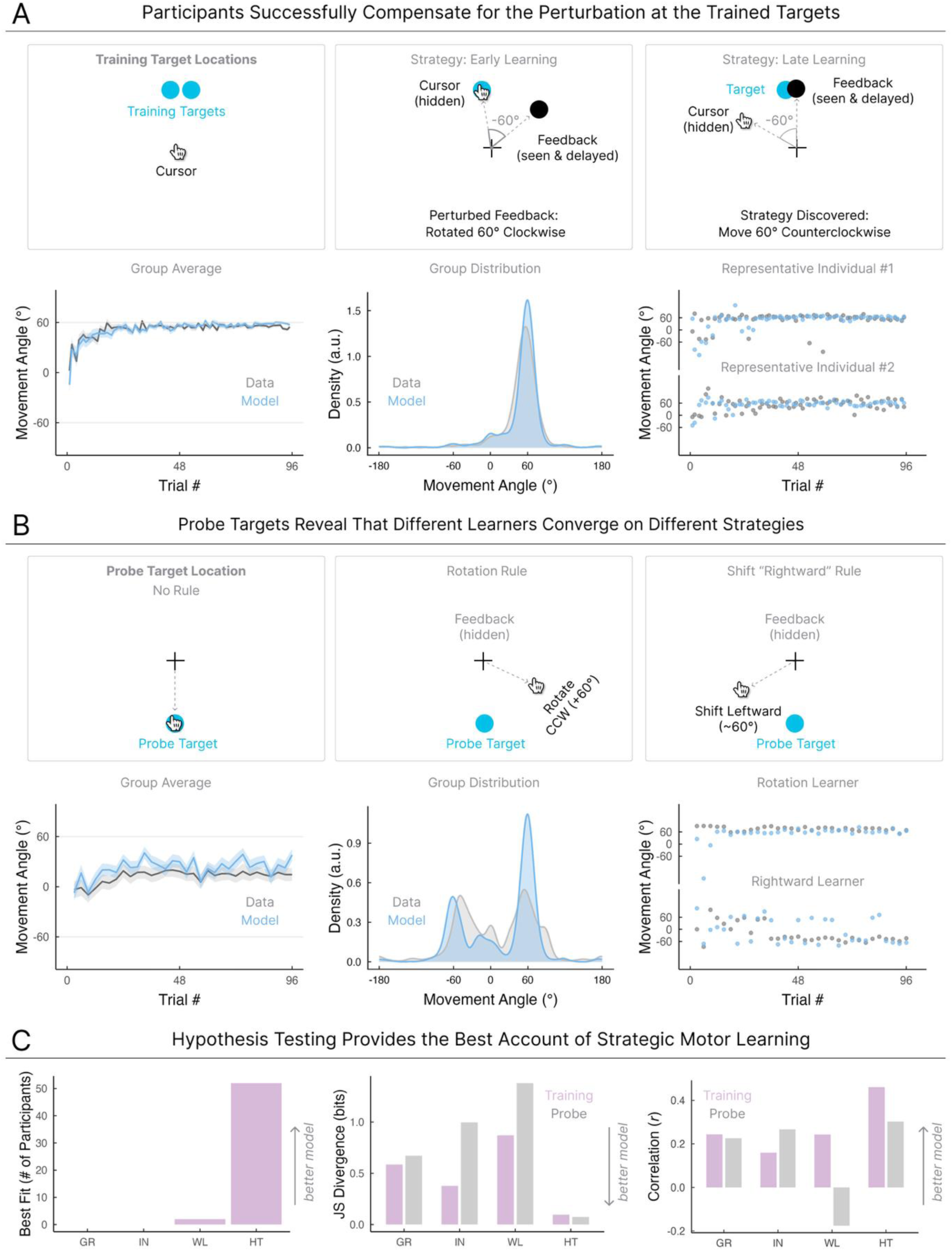
Hypothesis Testing Explains Why Different Learners Discover Different Solutions. **(A)** Experimental paradigm (top). Participants were instructed to make a straight, shooting movement through the target using their computer cursor, which remained hidden throughout the movement. A 60° clockwise visuomotor rotation was imposed, with rotated feedback (hollow black circle) presented as delayed endpoint feedback 800 ms after movement termination (perturbation was counterbalanced as counterclockwise or clockwise across participants), a manipulation known to minimize implicit motor learning, thereby isolating strategy discovery (Brudner et al., 2016). Participants learned to compensate for the perturbation at two closely spaced training targets. Group mean movement angles (solid line) ± 1 SEM (shaded) are shown (left), together with the distribution of movement angles (middle-left). Representative learning trajectories (right) from two participants who successfully compensated for the perturbation at the trained targets. **(B)** Probe trials, interleaved every three training trials, revealed whether participants adopted distinct motor strategies (top). If participants inferred a clockwise rotational rule, they should reach counterclockwise relative to the probe target; if they inferred a shift “rightward” rule, they should reach clockwise (“leftward”). Group mean movement angles (solid line) ± 1 SEM (shaded) are shown (left), together with the distribution of probe responses (middle-left). Representative learning trajectories from participants who converged on the rotational solution, rotating counterclockwise (top right), or the rightward-shift solution, shifting leftward (bottom right), illustrate how identical training at closely spaced targets led different learners to adopt distinct solutions. **(C)** Model comparison. Number of participants best fit by each model according to BIC (left). Jensen–Shannon distance between empirical and simulated movement-angle distributions for training (purple) and probe (grey) trial data (middle; lower values indicate better agreement). Pearson’s correlation between individuals’ empirical and simulated changepoints for training (purple) and probe (grey) (right; higher values indicate better agreement).

We first asked whether learning in this dataset exhibited the behavioral signatures of hypothesis testing. Participants successfully compensated for the perturbation at the trained targets (late learning = 54.3° ± 2.6°) and exhibited the two hallmarks of hypothesis testing with four prominent response modes centered near the correct solution (60°), the target (0°), a direction orthogonal to the targets (90°), and the sign-flip solution (−60°). Second, participants exhibited an ‘aha’ moment, the timing of which varied substantially across individuals (range: 2–52 training trials after perturbation onset). The presence of these signatures in an independent dataset suggests that they reflect general computational principles of strategy discovery rather than idiosyncratic features of a particular paradigm.

Having established these signatures, we next exploited the sensory ambiguity of the training regime to determine which hypothesis each learner ultimately adopted by interleaving no-feedback probe trials at a target 180° from the trained region. At this probe location, the two visuomotor rules make opposing predictions: If learners adopted a global rotation rule, they should compensate opposite the imposed perturbation (i.e., a +60° counterclockwise compensatory strategy). In contrast, if learners adopted a “shift rightward” rule, they should compensate in the same direction as the imposed perturbation (i.e., a −60° “leftward” compensatory strategy). The hypothesis testing account predicts that this ambiguity should lead *different* learners to commit to *different* hypotheses, producing a bimodal distribution of probe responses across individuals. By contrast, all alternative models, without additional assumptions, predict that learners converge on the same solution, yielding distributions near the rotational (+60°) solution.

Strikingly, the learners segregated into two distinct clusters: 52% of the participants adopted the global rotation rule (late probe movement angle = 55.0° ± 20.0°), whereas 48% adopted the shift-rightward rule (−31.4° ± 24.2°). Consistent with this bifurcation, simulations from the hypothesis testing model at the probe location were also multimodally distributed (Fig. 4B), with prominent peaks near +60° and −60°, corresponding to the rotational and shift rightward solutions. Thus, the hypothesis testing model captured the observation that different learners converged on different solutions to the same perturbation.

Quantitatively, the hypothesis testing model provided the best overall account of the data, outperforming all alternative models for every participant (54/54). It yielded the lowest Jensen–Shannon divergence for the distribution of movement angles during both training (0.10 bits; all alternative models > 0.20 bits) and probe trials (0.08 bits; all alternative models > 0.44 bits), indicating that it best captured the exploratory structure of participants’ behavior. The hypothesis testing model also provided the best account for the timing of the “aha” moment at both the training target (*r* = .46, p < .001, 95% CI [.2, .7]; alternative models *r* < .24) and probe target (Probe: *r* = .30, p = .03, 95% CI [-.4, .3]; alternative models *r* < .27). Taken together, these findings support a core prediction of hypothesis testing: when sensory evidence is consistent with multiple solutions, different learners can discover different solutions.

## Discussion

How do humans discover new motor strategies? Many of our most important motor behaviors—from a surgeon mastering an unfamiliar instrument to a patient relearning to move after injury—require us to first discover an effective movement before refining it through practice. Despite the ubiquity of this problem, the computational principles underlying motor strategy discovery remain unknown. A major obstacle has been the longstanding emphasis on gradual optimization—whether through error-based learning or win-stay lose-shift learning—supported by analyses that average behavior across learners and trials. By smoothing over rich behavioral variability, these approaches have obscured the very phenomena that define motor strategy discovery.

### Hypothesis Testing as a Computational Mechanism for Motor Strategy Discovery

Here, we developed the Motor Inference and Discovery (MIND) task, a novel behavioral paradigm that isolates strategy discovery by minimizing contributions from sensorimotor noise and other forms of motor learning. This approach revealed two defining signatures of strategy discovery: systematic exploration across multiple candidate solutions followed by an “aha” transition to a successful solution. Importantly, we show that a hypothesis testing model best explains these dynamics. The success of this model in accounting for the data across three experiments suggests that hypothesis testing provides a parsimonious account of how humans discover new strategies during motor learning.

The first hallmark of motor strategy discovery was the emergence of multiple discrete modes of exploratory behavior. This pattern is difficult to reconcile with gradual error correction models, which predict incremental convergence toward a single solution (Donchin et al., 2003; Smith et al., 2006; Thoroughman & Shadmehr, 2000), and is similarly inconsistent with win-stay–lose-shift and sudden-insight accounts (Cashaback et al., 2019; Townsend et al., 2026), which formalize pre-discovery behavior as random exploration around a previously successful action or the default strategy of reaching toward the target. In contrast, the hypothesis testing model naturally predicts such multimodal structure: each mode corresponds to a candidate visuomotor rule under evaluation. Moreover, when sensory evidence is ambiguous, the hypothesis testing model uniquely predicts why different learners may converge on different motor strategies.

A second hallmark of motor strategy discovery was the presence of an “aha” moment (quantified using a model-free change-point analysis that estimates abrupt shifts in the mean and variability of behavior). This pattern is again difficult to reconcile with error correction accounts, which predict gradual improvement rather than abrupt behavioral shifts. Although the presence of such change points is consistent with insight-based accounts specifically designed to capture them (Townsend et al. 2026), the hypothesis testing account explains *why* they emerge. That is, insight arises when evidence accumulates in favor of a particular visuomotor hypothesis, triggering a shift from exploring competing hypotheses to exploiting a successful solution.

Several important future directions emerge from the present work. First, we focused exclusively on the performance of successful learners; however, consistent with previous studies (Butcher et al., 2017; Cisneros et al., 2026; Tsay et al., 2023), many individuals failed to discover an effective strategy (see Methods). We speculate that these non-learners may still engage in hypothesis testing but fail because they lack relevant visuomotor hypotheses, assign weak priors to them, or evaluate them noisily—possibilities that future experiments can test. Second, the hypothesis space itself remains to be empirically defined. As in many Bayesian models, our current hypothesis space was specified based on plausible candidate rules (Lucas et al., 2015; Piantadosi et al., 2016). Systematically testing behavior across a richer set of perturbations should, in a more data-driven manner, reveal which motor rules learners consider and how they are structured. Third, the search process can be made more realistic (Lieder & Griffiths, 2019). As currently implemented, the hypothesis testing model assumes parallel updating of candidate hypotheses. Alternatively, the participants may operate in a more serial mode, selecting only a single hypothesis or a small subset of hypotheses for evaluation, and moving on to alternatives only when the observed outcomes fail to support the current candidate(s). Finally, the mechanism governing the transition from exploration to exploitation remains unclear: when do learners continue testing alternative hypotheses, and when do they transition to exploiting a particular solution. Characterizing this transition will be critical for developing a more complete mechanistic account of how learners search, discover, and, ultimately, commit to a motor strategy (see Fig. S5 for a comprehensive evaluation of alternative hypothesis testing accounts).

### Broader Implications of Hypothesis Testing for Motor Learning

Hypothesis Testing provides a new lens through which to reinterpret classic findings in motor learning. Consider savings—the phenomenon whereby participants relearn a perturbation more rapidly upon re-exposure. The computational basis of savings has long been debated, with proposed explanations including increased sensitivity of the implicit learning system to repeated errors (Herzfeld et al., 2014), rapid retrieval of a previously learned re-aiming strategy (Avraham et al., 2021), and the re-emergence of a latent motor memory as learners reacclimate to a familiar sensorimotor context (Heald et al., 2021). The hypothesis testing framework offers a complementary perspective: prior experience reshapes the prior distribution over candidate visuomotor hypotheses, increasing the likelihood that previously successful rules are tested first. Rather than searching the hypothesis space anew, learners can preferentially evaluate solutions that have succeeded before, accelerating convergence on the correct strategy.

Hypothesis testing may provide a common computational framework that cuts across multiple forms of motor learning (Krakauer et al., 2019; Tsay et al., 2024; Yadav & Duque, 2023). During strategic motor adaptation, learners generate and evaluate candidate visuomotor transformations—such as rotations, translations, or reflections—to compensate for perturbations to an existing movement (e.g., a golfer adjusting her swing angle to compensate for a crosswind). During de novo skill acquisition, by contrast, learners generate and evaluate novel motor repertoires (e.g., a golfer learning a new swing), compositionally combining their constituent elements into a successful motor solution (Tian et al., 2026). In both cases, learning proceeds through the generation, evaluation, and selection of competing hypotheses. What distinguishes adaptation from de novo skill acquisition may therefore not be the underlying computation, but the space of candidate solutions over which it operates.

Our behavioral and computational findings also motivate a broader perspective when considering the neural basis of motor learning. A long history of neurophysiological studies has shown that motor learning is accompanied by systematic changes in preparatory activity within the motor cortex (Sheahan et al., 2016; Sun et al., 2022; Vyas et al., 2018; Vyas, O’Shea, et al., 2020), changes that have largely been interpreted through the lens of gradual error-based learning. We posit that Hypothesis Testing may offer a more parsimonious explanation of these findings. Cognitive circuits upstream of motor cortex may generate, evaluate, and select candidate action–outcome rules, which in turn would provide structured inputs that reorganize preparatory activity in motor cortex as learners discover a successful strategy (Kang et al., 2026; Perich et al., 2018). Testing this account will require simultaneously measuring neural activity across the cognitive–motor hierarchy and identifying neural dynamics underlying multimodal exploration and abrupt “aha” moments as they unfold on single trials during motor strategy discovery (Vyas, Golub, et al., 2020).

## Methods

### Experiments

#### Participants

We recruited a total of 200 participants through Prolific, an online participant recruitment website (119 males, aged 25.4 ± 0.2 years; 177 right-handers). Eligibility criteria included: (i) age between 18 and 30, (ii) native English proficiency, (iii) a Prolific approval rating >97 (out of 100), and (iv) normal or corrected-to-normal vision. All participants provided informed consent in accordance with policies approved by UC Berkeley and Carnegie Mellon University’s Institutional Review Boards. Participants received monetary compensation for their time.

#### Apparatus

Participants completed our web-based reaching experiment on their own devices (trackpad or optical mouse) via an internet browser. Stimulus position and size were scaled based on the pixel units of the participants’ devices. All stimulus and task parameters reported below are expressed in physical units (cm) as rendered on a standard reference 14” display (3024 × 1964 resolution, 254 pixels per inch, device pixel ratio = 2, logical/CSS resolution 1512 × 870). Participant screen size was recorded to ensure that participants were able to visualize the entire task workspace.

#### General Task Procedure

A large white square (16.0 × 16.0 cm on the reference display) was presented on a black background that defined the workspace. A white cross (0.8 cm) marked the central start position, and a blue circle (0.5 cm in diameter) indicated the target. Targets appeared at a fixed radial distance of 4.3 cm from the start position, at one of eight locations along an invisible circle (15°, 60°, 105°, 150°, 195°, 240°, 285°, and 330°, where 0° corresponds to the rightmost, vertical-middle direction) (Fig. 1A).

At the start of each trial, the participant’s cursor automatically reset to the center of the screen. The cursor, displayed as a hand icon, remained visible throughout the trial, a manipulation known to minimize implicit motor learning (Wong et al, 2019). After 300 ms, a target appeared. The participant moved the cursor and clicked the location in the workspace they believed would cause the endpoint feedback—a white dot (0.4 cm in diameter)—to appear on the target (Fig. 1A). A standard “click” sound confirmed their selection. Endpoint feedback was displayed immediately after the click and remained visible, along with the mouse cursor and target, for 2000 ms. The feedback appeared at the selected location (superimposed on the cursor) on veridical feedback trials and at a different location on perturbation trials (see below). If the perturbation caused the feedback to fall outside the workspace, no cursor was shown. Instead, a red “Out of Bounds” message appeared at the center of the screen for 2000 ms. Following the feedback interval, the feedback and target were removed, and the cursor reappeared at the center start position, marking the beginning of the next trial. The lower-right of the screen displayed the total number of trials along with a running trial count, allowing participants to monitor their progress.

The experiment consisted of two phases (160 trials total): a familiarization phase (8 trials) and a perturbation phase (152 trials). During the familiarization phase, endpoint feedback was veridical—the white dot appeared exactly at the cursor location at the time of the participant’s click. In the perturbation phase, feedback was systematically shifted relative to the participant’s click location. For Experiment 1, the perturbation was a 60° rotation (N = 50 Counterclockwise, N = 50 Clockwise). For Experiment 2, the perturbation was a diagonal mirror reflection (N = 100; reflection across a 45° axis crossing upper-left quadrant). Targets were presented in a pseudorandom sequence that avoided immediate repetitions and ensured that all targets were sampled once every eight trials (i.e., movement cycle). This sequence was consistent across participants to reduce variability that might be idiosyncratic to specific targets. Experiment 3 used a delayed-feedback task in which participants made rapid center-out reaching movements to counteract a visuomotor rotation. Endpoint feedback was presented 1000 ms after movement completion and remained visible for 500 ms, a manipulation shown to minimize implicit motor learning. For complete experimental details, see Ding et al. (2026).

#### Instructions

Before beginning the experiment, participants were given a simple directive: “Your goal is to land the white dot on the blue target.” Prior to the familiarization phase, they were told, “The white dot will land wherever you click,” and instructed to, “click directly on the blue target.” Before the perturbation phase, participants were informed, “We’ve changed the relationship between where you click and where the white dot appears. The white dot will no longer appear exactly where you click. This new relationship is consistent across all target locations. Your task is to figure out where to click to make the white dot land on the blue target.”

To ensure task comprehension, we embedded two multiple-choice instruction checks to verify their understanding of (1) the goal of the experiment (to identify where to click to land the feedback on the blue target) and (2) that there was a consistent transformation being applied to their movement decisions and feedback. These checks were administered after the instructions and before the movement of that trial. The first attention check occurred immediately following the task instructions about the perturbation (trial 9), and the second occurred three trials into the perturbation phase, after participants received a reminder of the task instructions (trial 12).

### Data Analysis

#### General Analysis Procedure

All analyses were conducted in R. We focused on participants’ click (i.e., choice) locations, analyzed in both Cartesian coordinates (X, Y) and polar coordinates (movement angle, radial distance relative to the target). Unless otherwise specified, data were analyzed across predefined phases: Baseline (trials 1-8), early learning (trials 9 – 24), and late learning (trials 145–160). For experiment 3, analyses were only performed on the perturbation phase of learning. Early learning was defined as the first 24 trials of learning (1-24; 8 cycles), and late learning as the last 24 trials (73-96; 8 cycles). When necessary for density visualization and statistical analyses, movement angles were converted to radians and analyzed using circular statistics to account for the periodic nature of angular data (Circular Statistics [R package circular version 0.5-2], 2025). For example, probability density of movement angles was estimated using a von Mises kernel density estimator, which accounts for the circular nature of angular data. Additionally, circular statistics were used to characterize movement angles and variability; paired t-tests were still used to assess statistical changes between early and late adaptation across participants.

#### Learner classification

Learners were defined based on performance during late learning. For each participant, we computed the mean absolute angular error during this phase, where error was defined as the angular distance between the target-specific solution and the movement angle selected on a given trial. We then fit a two-component Gaussian mixture model (R package: *mclust*) to these values across participants (Chassagnol et al., 2023). Participants assigned to the lower-error component were classified as learners, whereas those assigned to the higher-error component were classified as non-learners. Further checks were made to ensure the mean hand angles during late learning were different between groups (non-learners’ late-learning hand angle from target, where solution is at +60°: Experiment 1 = 6.4° ± 0.01°; Experiment 2 = 21.9° ± 0.1°). Non-learners were excluded from subsequent analyses to focus on participants who successfully discovered a viable strategy. Applying this criterion resulted in 50 learners out of 100 participants (50% learner rate) in the rotation group and 44 learners out of 100 participants in the mirror reflection group (44% learner rate). These learner rates are comparable to those reported in previous studies of strategic motor learning (Cisneros et al., 2026; Tsay et al., 2023), suggesting that failure to discover an effective strategy is a robust feature of these tasks rather than an idiosyncrasy of the present experiments. Understanding why some individuals fail to discover an effective strategy remains an important question and will be the focus of future work.

#### Gaussian mixture model

To identify common systematic patterns associated with strategy discovery, we fit Gaussian mixture models (R package: *mclust*). Analyses were performed on the first 80 perturbation trials to obtain stable estimates of the underlying movement modes. The optimal number of modes was selected using the Bayesian Information Criterion (BIC). To quantify evidence for multimodality, we report the BIC difference (ΔBIC) between the best-fitting model with more than one component and the best-fitting single-component model. Lower ΔBIC values indicate stronger evidence for multimodality. For visualization, cluster means were transformed from Cartesian coordinates to preserve their angular positions while normalizing their radial distances to match those of the target locations.

#### Change-point Analysis

To quantify the “aha” moment associated with strategy discovery, we applied the nonparametric, model-free Pruned Exact Linear Time (PELT) algorithm (R package: changepoint; function: cpt.meanvar) to each participant’s angular movement error. This method detects abrupt changes in the mean and variance of the error distribution. We imposed a minimum segment length of a training cycle, preventing changepoints from occurring within the first cycle of perturbed trials (eight trials for Experiments 1 & 2, two trials for Experiment 3). The first detected changepoint was taken as the time of strategy discovery. A changepoint was identified for all participants in Experiment 1, all but one participant in Experiment 2, and all but one participant in Experiment 3. Participants for whom no changepoint was detected were excluded from changepoint analyses. For Experiment 3, a changepoint was detected separately for training and probe trials, and the training results are reported in the behavioral results section.

#### Clustering participants

Participant-level clustering was performed on the data from Experiment 3 using an approach adapted from Ding et al. (2026). Pairwise dissimilarities between participants’ learning trajectories were quantified using dynamic time warping (Giorgino, 2009), with cosine distance as the similarity metric. The resulting pairwise distance matrix served as the input for hierarchical agglomerative clustering using Ward’s minimum-variance method. The optimal number of clusters, evaluated across solutions ranging from 2 to 30, was determined using both the average silhouette score and within-cluster sum of squares.

#### Computational Modeling

To distinguish among competing accounts of strategy discovery, we fit participants’ movement choices to four candidate models spanning a range of learning mechanisms.

#### Hypothesis Testing Model

This model assumes that learners seek to infer the hidden visuomotor rule linking their actions to visual feedback. To do so, they maintain a set of candidate action–outcome rules— including rotations, reflections, gains, and translations. We formalized this hypothesis space as 29 discrete hypotheses (Table S1), chosen to capture a range of plausible explanations that participants might entertain when confronted with the perturbation. The set of candidate hypotheses, *H*, was denoted as:

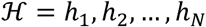

*N* denotes the total number of hypotheses. At the start of learning, all hypotheses were assumed to be equally plausible and were therefore assigned a uniform prior probability:

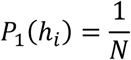

On each trial (t), learners selected a hypothesis according to their current belief distribution *P_t_*(ℎ*_i_*) and generated an action consistent with that hypothesis.

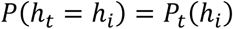

Once a hypothesis was selected, the learner generated the movement expected to bring the feedback cursor to the target location. The target location on trial *t* is denoted (*g_t_*). Actions (*x_t_*) were generated by applying the inverse of the selected hypothesis to the target location, producing the action expected to bring the feedback cursor to the target under the learner’s current hypothesis. Motor execution was assumed to be subject to Gaussian motor noise, 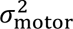.

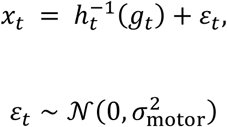

After observing the resulting sensory outcome, *y_t_*, the learner evaluated how well the predicted outcome for each candidate hypothesis, *ŷ_i,t_*, accounted for the observed feedback. Observed outcomes were assumed to be subject to Gaussian observation noise, 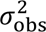.

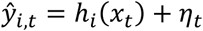

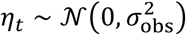

Under this assumption, the likelihood of the observed feedback under hypothesis (ℎ*_i_*) is given by

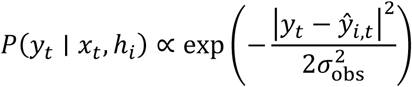

such that hypotheses that generated feedback predictions *ŷ_i_*,*_t_* closer to the observed feedback *y_t_* received greater support. Beliefs were then updated according to Bayes’ rule,

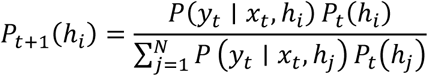

Consequently, hypotheses that better explained the observed feedback became increasingly probable, whereas poorly supported hypotheses were progressively eliminated. As evidence accumulated across trials, the posterior distribution concentrated on a small subset of candidate hypotheses, ultimately enabling convergence on a successful strategy. The model contains two free parameters: a motor noise parameter (*σ*_motor_) and an observation noise parameter (*σ*_obs_).

#### Gradual Error Reduction

This model represents a passive error-based learning process in which the action policy gradually shifts toward a successful solution as evidence accumulates. Although many implementations are possible (Albert et al., 2022; Sugiyama et al., 2023), we designed this model to parallel the hypothesis testing model closely.

Specifically, the model assumes that learners maintain beliefs over a reduced hypothesis space consisting of only two possibilities: no perturbation (ℎ_null_) and the true perturbation (ℎ_true_). Beliefs are updated according to the same Bayesian update rule described above. However, unlike the hypothesis testing model, learners do not sample individual hypotheses. Instead, actions are generated according to a posterior-weighted average of the two candidate policies:

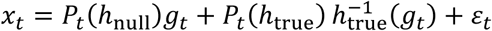

Consequently, as evidence accumulates in favor of the true perturbation, the action policy gradually shifts from aiming directly at the target toward full compensation for the imposed perturbation. This was enforced by increasing the prior probability of the null hypothesis *P*_0_(ℎ_null_) and decreasing the prior of the true hypothesis, *P*_0_(ℎ_true_). This model predicts smooth and continuous error reduction over time. The model contains two free parameters: a motor noise parameter (*σ*_motor_) and an observation noise parameter (*_σ_*obs).

#### Insight Model

The Insight model assumes that learners initially behave as though no perturbation is present and then abruptly discover the correct solution. The hypothesis space consists of only two possibilities: no perturbation (ℎ*_null_*) and the true perturbation (ℎ*_true_*). Prior to the moment of insight, defined by a changepoint (*τ*), actions are generated according to the null hypothesis, reflecting the assumption that learners have not yet identified the underlying perturbation:

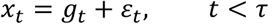

To allow for flexibility across experiments, this parameter, (*τ*), was fitted between values of 0 and 1, and transformed by multiplying this value by the total number of trials within the experiment. The fitted parameter serves as the fraction of total time before insight occurs. Following insight, actions are generated according to a weighted combination of the null policy and the fully compensatory policy,

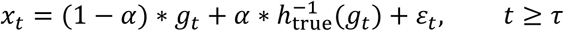

where(*α* ∈ [0,1]) determines the extent to which the inferred rule is applied. When (*α* = 0), behavior remains unchanged from the null policy. When (*α* = 1), the learner fully compensates for the perturbation; intermediate values correspond to partial implementation of the discovered rule (Townsend et al., 2025). The model contains three free parameters: an insight-time parameter (*_τ_*), an implementation parameter (*α*), and a motor noise parameter (*σ*_motor_).

#### Win-Stay, Lose-Shift

The Win-Stay, Lose-Shift model assumes that learners refine their aim based on the success or failure of previous movements rather than by maintaining explicit beliefs about candidate perturbations (Cashaback et al., 2019; Therrien et al., 2016; van Mastrigt et al., 2021). The model operates directly in Cartesian screen coordinates (x, y) and maintains an independent cached policy for each target, such that the history of successes and failures at one target provides no information about how to act at another.

For a given target, the model maintains a cached aim location *μ_t_* = (*μ_x,t_*, *μ_y,t_*) and movement variability *σ_t_*. The movement produced on trial *t* is drawn as:

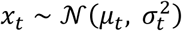

where variability is assumed independent and equal along the x- and y-axes. Before any feedback has been observed, the cached aiming location is initialized at the target location, and movement variability reflects motor execution noise, 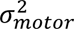.

Following movement execution, success is determined by the Euclidean distance, *δ_t_*, between the (perturbed) feedback cursor position and the target, relative to a fixed reward threshold, *τ*. This reward threshold corresponds to the maximum size of the region for which a movement is considered successful. If the movement is successful, the executed action is cached as the new aiming angle, and subsequent variability is reduced to motor noise alone (“win-stay”). If the movement is unsuccessful, the cached aiming angle remains unchanged but variability increases, promoting broader exploration around the previously cached action (“lose-shift”):

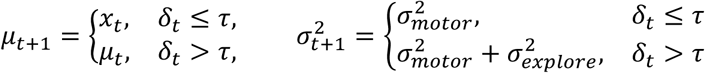

Exploration noise was represented as a scaled version of motor noise:

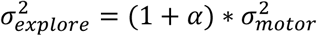

ensuring that exploratory noise exceeded motor noise. The model thus contains three free parameters: motor noise (*σ_motor_*), the exploration scaling parameter (*α*), and the reward threshold (*τ*).

#### Model Fitting, Comparison, and Validation

Model parameters were estimated separately for each participant using maximum likelihood estimation implemented with Python’s *scipy.optimize.minimize* function. To reduce the risk of convergence to local minima, each model was fit 30 times from randomly initialized starting values, and the solution with the lowest negative log-likelihood was retained. Model comparisons were performed using the Bayesian Information Criterion (BIC). Parameter ranges were chosen to span a broad range of plausible behavioral signatures, capturing the different ways each model could behave while remaining within realistic bounds of motor and sensorimotor noise (Table S2). To further evaluate model performance, we simulated behavior using each participant’s best-fitting parameter estimates (Fig. S4). For each participant and model, we generated 1,000 simulated datasets and quantified the similarity between each simulation and the empirical data using the circular sum-of-squared-errors (SSE). For posterior predictive checks across models, we selected the simulation with the median SSE relative to the empirical data. For Experiment 3, we selected the simulation with the lowest SSE to determine whether any model simulation could reproduce the signatures observed in the probe data.

To validate how well each model captured key behavioral signatures of learning, we computed three metrics: (i), we quantified the similarity between empirical and simulated early-learning movement distributions using Jensen-Shannon divergence (Lin, 1991) across individuals (ii) the changepoint correlation, defined as the Pearson correlation between the observed and simulated changepoint for each individual and (iii) correlations between empirical and simulated individual movement variability (standard deviation of movement angle). Together, these analyses provided a stringent test of each model’s ability to reproduce both the distributional structure of systematic exploration and the timing of the “aha” moment.

We further conducted parameter and model recovery analyses (Fig. S2) (Wilson & Collins, 2019). For parameter recovery, we generated synthetic datasets from each model using 100 randomly sampled parameter combinations spanning the fitted parameter space. The models were then refit to these datasets, and recovery was quantified by comparing the recovered parameter estimates with their known generating values. For model recovery, we generated 100 synthetic datasets from each candidate model and evaluated whether the true generating model was correctly identified using BIC. All candidate models were reliably recovered, with particularly strong discrimination between the hypothesis testing model and alternative models. Parameter recovery was similarly robust across all models. Together, these analyses demonstrate that our fitting procedure reliably recovers both the model identity and the underlying parameters, thereby supporting the robustness and interpretability of our modeling framework.

## Acknowledgements

This material is based upon work supported by the National Science Foundation Graduate Research Fellowship Program under Grant No(s) (NSF DGE2631988). Any opinions, findings, and conclusions or recommendations expressed in this material are those of the author(s) and do not necessarily reflect the views of the National Science Foundation. Additionally, this work is funded by NSF Grant No. 2545300 awarded to Jonathan Tsay. We thank Xaq Pitkow for insightful discussions that helped strengthen this work.

## Supplemental Materials

**Figure S1.**
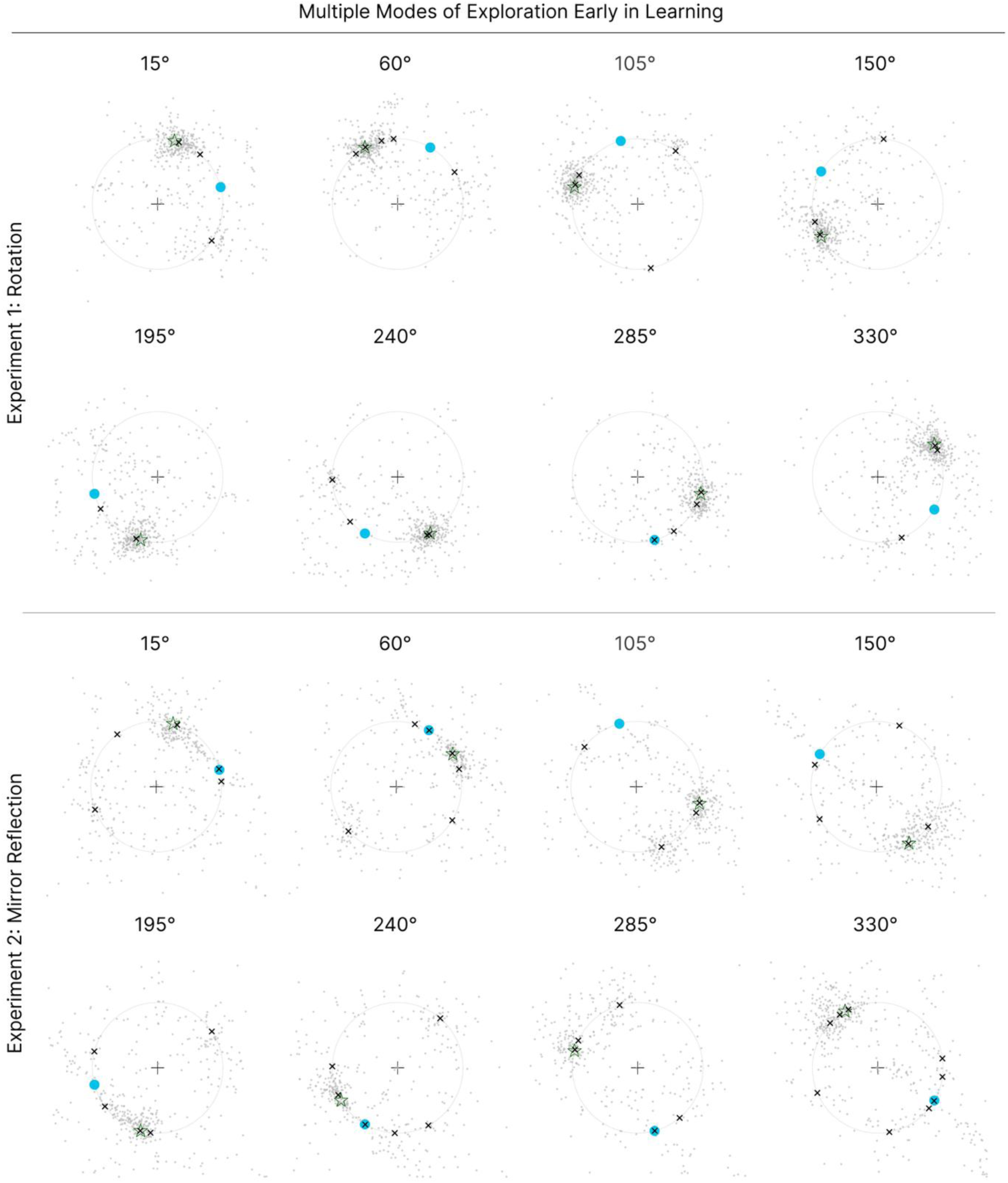
Multiple modes of exploration during early learning. Gaussian mixture model (GMM) clustering of motor responses for each target location (columns) and experiment (rows) during early learning (first 80 perturbed trials). Each gray point represents the reported movement location from one participant on a single trial. The blue circle denotes the target location, the green star the motor solution, and black ×’s the angular peaks identified by the clustering algorithm.

**Figure S2.**
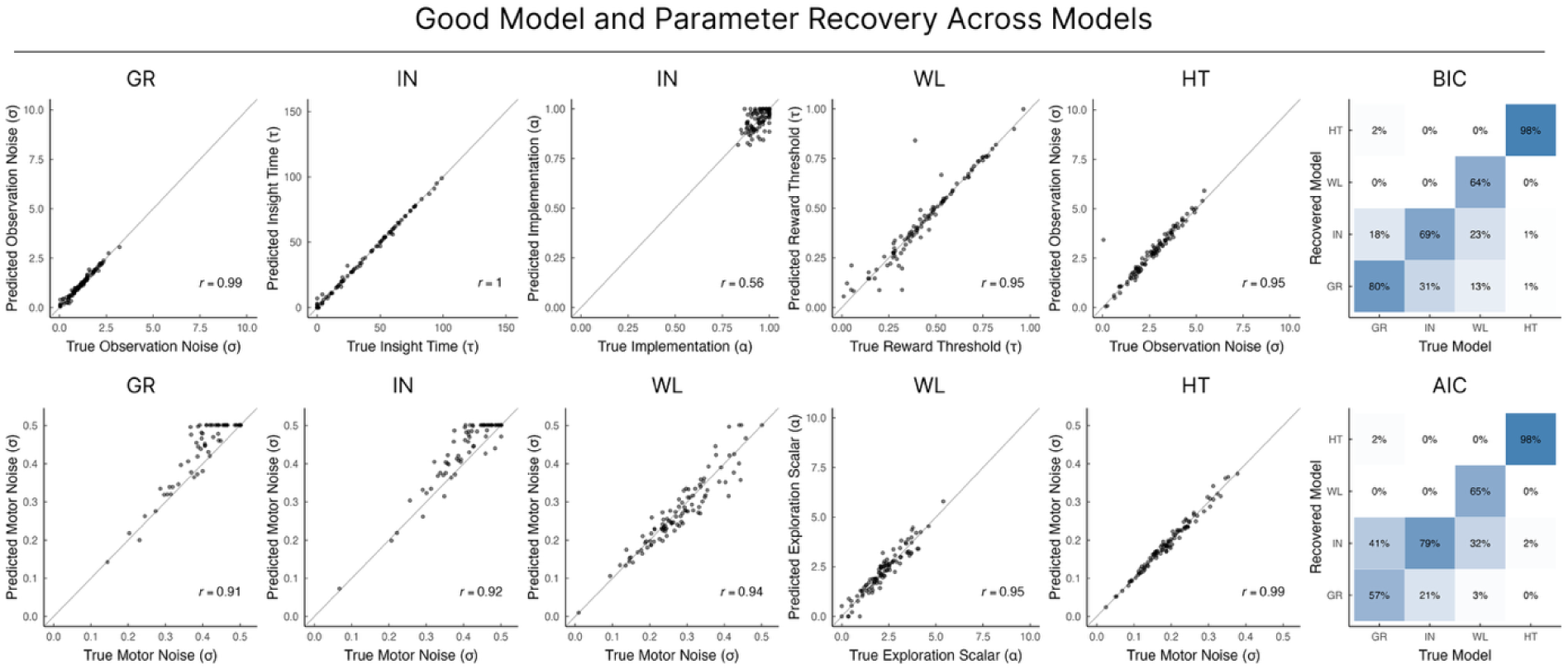
Model and parameter recovery. Parameter recovery for the four comparison models, showing the correspondence between true and recovered parameter values. Each point represents parameter estimates from one simulated dataset. The gray identity line indicates ideal recovery, and *r* denotes the correlation between true and recovered parameter values. Model recovery confusion matrix (rightmost column) using BIC (top) and AIC (bottom). Data were simulated from each candidate model (columns) and refit with all four models. Cell values indicate the percentage of 100 simulated datasets generated by each model that were best fit by each recovered model.

**Figure S3.**
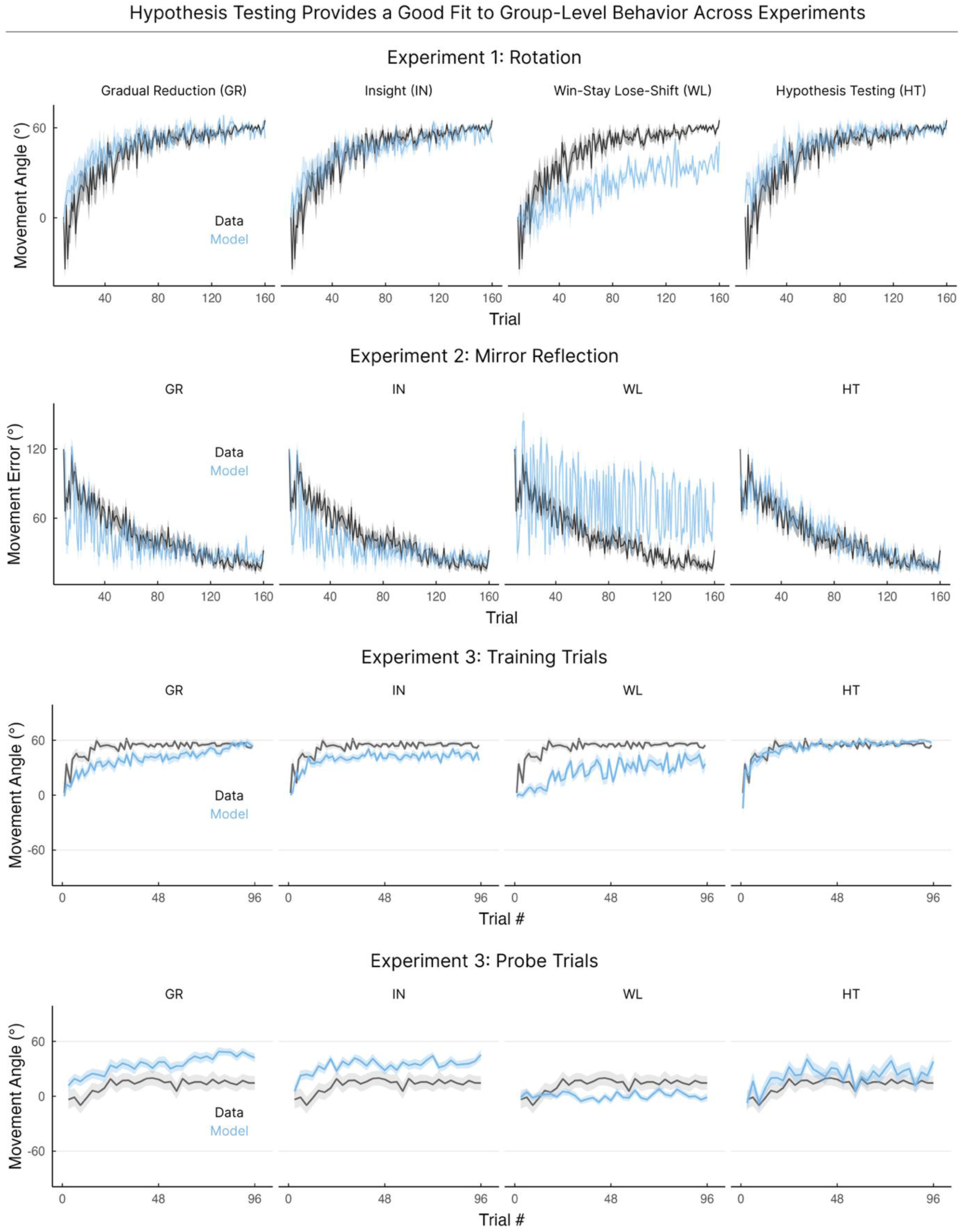
Group learning trajectories for Experiments 1-3. Empirical behavior is shown in black, with corresponding model simulations of the group-averaged learning trajectories shown in blue. Columns correspond to the competing models. For Experiment 2, movement error is plotted to summarize learning across all target locations. For Experiment 3, training data is shown above, and probe data below.

**Figure S4.**
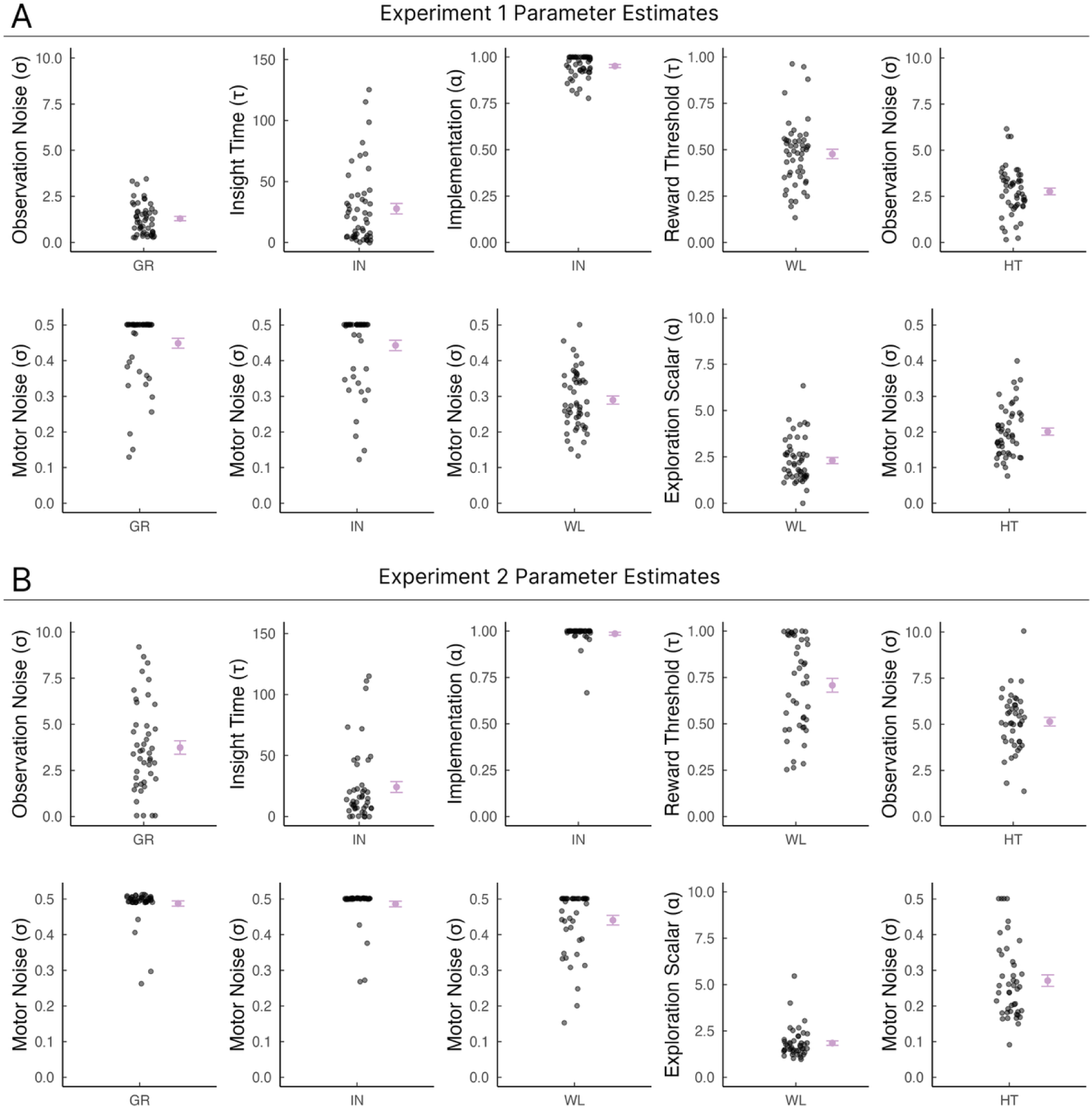
Parameter estimates for Experiments 1 (A) and 2 (B). Best-fitting parameter estimates for each of the four candidate models. The top row shows the primary model parameters, and the bottom row shows estimated motor noise. Each black point represents one participant’s best-fitting parameter estimate; purple points and error bars indicate the group means ± SEM.

**Figure S5.**
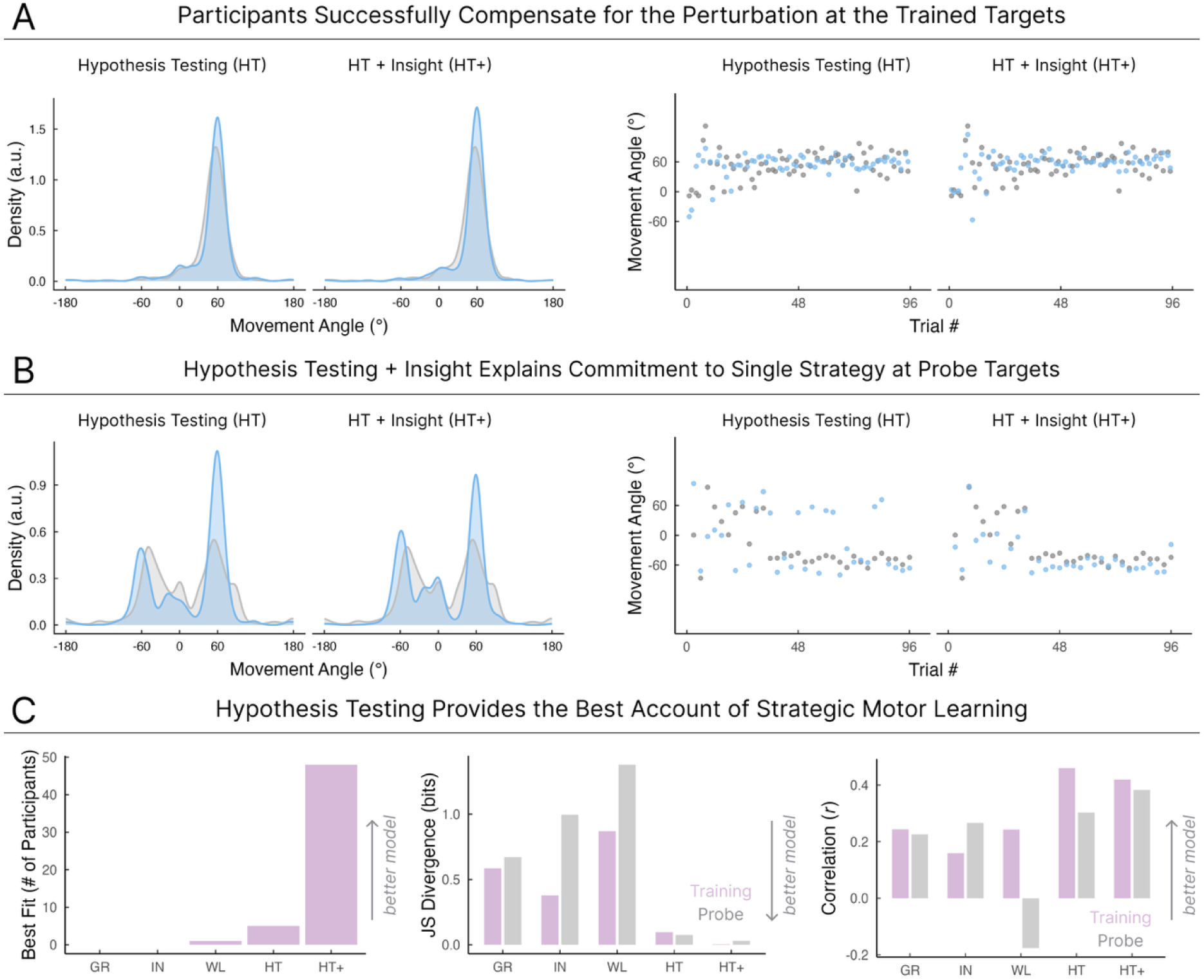
A hybrid model that combines Hypothesis Testing and Insight best explains behavior in Experiment 3. In post hoc analyses, we considered a variant of the Hypothesis Testing model in which learners commit to a single hypothesis after a sudden moment of insight, potentially reflecting a decision threshold beyond which continued exploration is no longer necessary (HT + Insight, or HT+). We compared this variant, which included an additional changepoint parameter, with the basic Hypothesis Testing (HT) model. The left column shows results for the basic HT model, and the right column shows results for HT+. **(A)** Left shows distribution of movement angles during early training trials. Trial-by-trial movement angle data for a representative participant are shown in grey and predicted model results are shown in blue. **(B)** Same as A but for probe trials. **(C)** Number of participants best fit by each model according to BIC (left). Jensen–Shannon divergence between group-level empirical and simulated early movement angle distributions during training (purple) and probe (grey) (middle; lower values indicate better agreement). Pearson’s correlation between individuals’ empirical and simulated changepoints (right; higher values indicate better agreement).

**Table S1.** Affine transformations included in the hypothesis space of the Hypothesis Testing model, expressed in matrix form.

| Rule | Matrix | Rule | Matrix | Rule | Matrix |
| --- | --- | --- | --- | --- | --- |
| Reflection (0°) | $\begin{bmatrix} 1 & 0 \\ 0 & -1 \end{bmatrix} \begin{bmatrix} x \\ y \end{bmatrix}$ | Shift Leftward | $\begin{cases} \text{Rotation}(+60^\circ) & \text{if } 45^\circ \leq \theta < 225^\circ \\ \text{Rotation}(-60^\circ) & \text{otherwise} \end{cases}$ | Translation (0.2, -0.2) | $\begin{bmatrix} 1 & 0 & 0.2 \\ 0 & 1 & -0.2 \\ 0 & 0 & 1 \end{bmatrix} \begin{bmatrix} x \\ y \\ 1 \end{bmatrix}$ |
| Reflection (45°) | $\begin{bmatrix} 0 & 1 \\ 1 & 0 \end{bmatrix} \begin{bmatrix} x \\ y \end{bmatrix}$ | Shift Downward | $\begin{cases} \text{Rotation}(+60^\circ) & \text{if } 135^\circ \leq \theta < 315^\circ \\ \text{Rotation}(-60^\circ) & \text{otherwise} \end{cases}$ | Translation (-0.2, 0.2) | $\begin{bmatrix} 1 & 0 & -0.2 \\ 0 & 1 & 0.2 \\ 0 & 0 & 1 \end{bmatrix} \begin{bmatrix} x \\ y \\ 1 \end{bmatrix}$ |
| Reflection (90°) | $\begin{bmatrix} -1 & 0 \\ 0 & 1 \end{bmatrix} \begin{bmatrix} x \\ y \end{bmatrix}$ | Shift Rightward | $\begin{cases} \text{Rotation}(+60^\circ) & \text{if } \theta \geq 225^\circ \text{ or } \theta < 45^\circ \\ \text{Rotation}(-60^\circ) & \text{otherwise} \end{cases}$ | Translation (-0.4, 0.4) | $\begin{bmatrix} 1 & 0 & -0.4 \\ 0 & 1 & 0.4 \\ 0 & 0 & 1 \end{bmatrix} \begin{bmatrix} x \\ y \\ 1 \end{bmatrix}$ |
| Reflection (135°) | $\begin{bmatrix} 0 & -1 \\ -1 & 0 \end{bmatrix} \begin{bmatrix} x \\ y \end{bmatrix}$ | Shift Upward | $\begin{cases} \text{Rotation}(+60^\circ) & \text{if } \theta \geq 315^\circ \text{ or } \theta < 135^\circ \\ \text{Rotation}(-60^\circ) & \text{otherwise} \end{cases}$ | Translation (0.5, -0.5) | $\begin{bmatrix} 1 & 0 & 0.5 \\ 0 & 1 & -0.5 \\ 0 & 0 & 1 \end{bmatrix} \begin{bmatrix} x \\ y \\ 1 \end{bmatrix}$ |
| Rotation (-180°) | $\begin{bmatrix} -1 & 0 \\ 0 & -1 \end{bmatrix} \begin{bmatrix} x \\ y \end{bmatrix}$ | Gain (0.5) | $\begin{bmatrix} 0.5 & 0 \\ 0 & 0.5 \end{bmatrix} \begin{bmatrix} x \\ y \end{bmatrix}$ | Translation (-0.5, 0.5) | $\begin{bmatrix} 1 & 0 & -0.5 \\ 0 & 1 & 0.5 \\ 0 & 0 & 1 \end{bmatrix} \begin{bmatrix} x \\ y \\ 1 \end{bmatrix}$ |
| Rotation (-170°) | $\begin{bmatrix} \cos(-170^\circ) & -\sin(-170^\circ) \\ \sin(-170^\circ) & \cos(-170^\circ) \end{bmatrix} \begin{bmatrix} x \\ y \end{bmatrix}$ | Gain (1.5) | $\begin{bmatrix} 1.5 & 0 \\ 0 & 1.5 \end{bmatrix} \begin{bmatrix} x \\ y \end{bmatrix}$ | Translation (-0.6, 0.6) | $\begin{bmatrix} 1 & 0 & -0.6 \\ 0 & 1 & 0.6 \\ 0 & 0 & 1 \end{bmatrix} \begin{bmatrix} x \\ y \\ 1 \end{bmatrix}$ |
| Rotation (-120°) | $\begin{bmatrix} \cos(-120^\circ) & -\sin(-120^\circ) \\ \sin(-120^\circ) & \cos(-120^\circ) \end{bmatrix} \begin{bmatrix} x \\ y \end{bmatrix}$ | | | Translation (-0.7, 0.7) | $\begin{bmatrix} 1 & 0 & -0.7 \\ 0 & 1 & 0.7 \\ 0 & 0 & 1 \end{bmatrix} \begin{bmatrix} x \\ y \\ 1 \end{bmatrix}$ |
| Rotation (-60°) | $\begin{bmatrix} \cos(-60^\circ) & -\sin(-60^\circ) \\ \sin(-60^\circ) & \cos(-60^\circ) \end{bmatrix} \begin{bmatrix} x \\ y \end{bmatrix}$ | | | Translation (0.8, -0.8) | $\begin{bmatrix} 1 & 0 & 0.8 \\ 0 & 1 & -0.8 \\ 0 & 0 & 1 \end{bmatrix} \begin{bmatrix} x \\ y \\ 1 \end{bmatrix}$ |
| Identity (0°) | $\begin{bmatrix} 1 & 0 \\ 0 & 1 \end{bmatrix} \begin{bmatrix} x \\ y \end{bmatrix}$ | | | Translation (1.2, -1.2) | $\begin{bmatrix} 1 & 0 & 1.2 \\ 0 & 1 & -1.2 \\ 0 & 0 & 1 \end{bmatrix} \begin{bmatrix} x \\ y \\ 1 \end{bmatrix}$ |
| Rotation (60°) | $\begin{bmatrix} \cos 60^\circ & -\sin 60^\circ \\ \sin 60^\circ & \cos 60^\circ \end{bmatrix} \begin{bmatrix} x \\ y \end{bmatrix}$ | | | Translation (-1.3, 1.3) | $\begin{bmatrix} 1 & 0 & -1.3 \\ 0 & 1 & 1.3 \\ 0 & 0 & 1 \end{bmatrix} \begin{bmatrix} x \\ y \\ 1 \end{bmatrix}$ |
| Rotation (120°) | $\begin{bmatrix} \cos 120^\circ & -\sin 120^\circ \\ \sin 120^\circ & \cos 120^\circ \end{bmatrix} \begin{bmatrix} x \\ y \end{bmatrix}$ | | | | |
| Rotation (170°) | $\begin{bmatrix} \cos 170^\circ & -\sin 170^\circ \\ \sin 170^\circ & \cos 170^\circ \end{bmatrix} \begin{bmatrix} x \\ y \end{bmatrix}$ | | | | |
| Rotation (180°) | $\begin{bmatrix} -1 & 0 \\ 0 & -1 \end{bmatrix} \begin{bmatrix} x \\ y \end{bmatrix}$ | | | | |

**Table S2.** Parameter ranges across models were selected to capture a broad range of behavioral patterns.

| **Model** | **Parameter** | **Min** | **Max** |
| --- | --- | --- | --- |
| **GR** | Observation Noise ($\sigma$) | 0 | 5 |
| **GR** | Motor Noise ($\sigma$) | 0 | 0.5 |
| **IN** | Insight Time ($\tau$) | 0 | 1 |
| **IN** | Implementation ($\alpha$) | 0 | 1 |
| **IN** | Motor Noise ($\sigma$) | 0 | 0.5 |
| **WL** | Reward Threshold ($\tau$) | 0.01 | 1 |
| **WL** | Exploration Scalar ($\alpha$) | 0 | 10 |
| **WL** | Motor Noise ($\sigma$) | 0 | 0.5 |
| **HT** | Observation Noise ($\sigma$) | 0 | 10 |
| **HT** | Motor Noise ($\sigma$) | 0 | 0.5 |

